# 17α-estradiol, a lifespan-extending compound, attenuates liver fibrosis by modulating collagen turnover rates in male mice

**DOI:** 10.1101/2022.06.16.496423

**Authors:** Samim Ali Mondal, Roshini Sathiaseelan, Shivani N. Mann, Maria Kamal, Wenyi Luo, Tatiana D. Saccon, José V.V. Isola, Frederick F. Peelor, Tiangang Li, Willard M. Freeman, Benjamin F. Miller, Michael B. Stout

**Author notes:** **Correspondence To:** Michael B. Stout, Oklahoma Medical Research Foundation, 825 NE 13th Street, Chapman S212, Oklahoma City, OK 73104. **Email Addresses of Authors:** Samim Ali Mondal Roshini Sathiaseelan Shivani N. Mann Maria Kamal Wenyi Luo Tatiana D. Saccon José V. V. Isola Frederick F. Peelor III Tiangang Li Willard M. Freeman Benjamin F. Miller.

## Abstract

**Background:** Estrogen signaling is protective against chronic liver diseases, although men and a subset of women are contraindicated for chronic treatment with 17β-estradiol (17β-E2) or combination hormone replacement therapies. We sought to determine if 17α-estradiol (17α-E2), a naturally-occurring diastereomer of 17β-E2, could attenuate liver fibrosis.

**Methods:** We evaluated the effects of 17α-E2 treatment on collagen synthesis and degradation rates using tracer-based labeling approaches in male mice subjected to carbon tetrachloride (CCl_4_)-induced liver fibrosis. We also assessed the effects of 17α-E2 on markers of hepatic stellate cell (HSC) activation, collagen crosslinking, collagen degradation, and liver macrophage content and polarity.

**Findings:** We found that 17α-E2 significantly reduced collagen synthesis rates and increased collagen degradation rates, which was mirrored by declines in transforming growth factor β1 (TGF-β1) and lysyl oxidase-like 2 (LOXL2) protein content in liver. These improvements were associated with increased matrix metalloproteinase 2 (MMP2) activity and suppressed stearoyl-coenzyme A desaturase 1 (SCD1) protein levels, the latter of which has been linked to the resolution of liver fibrosis. We also found that 17α-E2 increased liver fetuin-A protein, a strong inhibitor of TGF-β1 signaling, and reduced pro-inflammatory macrophage activation and cytokines expression in the liver.

**Interpretation:** We conclude that 17α-E2 reduces fibrotic burden by suppressing HSC activation and enhancing collagen degradation mechanisms. Future studies will be needed to determine if 17α-E2 acts directly in hepatocytes, HSCs, and/or immune cells to elicit these benefits.

**Funding:** This work was supported by the US National Institutes of Health and US Department of Veterans Affairs.

**RESEARCH IN CONTEXT:** *Evidence before this study:* The prevalence and severity of chronic liver diseases are greater in men than women and men are twice as likely to die from chronic liver diseases. However, the prevalence and severity of nonalcoholic fatty liver disease (NAFLD), nonalcoholic steatohepatitis (NASH), and liver fibrosis becomes comparable between the sexes following menopause, particularly when hormone replacement therapies (HRT) are not initiated. These observations suggest that estrogen signaling is protective against liver disease onset and progression, which is supported by studies in rodents demonstrating that 17β-estradiol (17β-E2) ameliorates hepatic steatosis and fibrogenesis. However, chronic administration of 17β-E2 or combination HRTs are unrealistic in men due to feminization and increased risk for stroke and prostate cancer, and a subset of the female population are also at an increased risk for breast cancer and cardiovascular events when on HRTs. Therefore, we have begun exploring the therapeutic potential of 17α-estradiol (17α-E2), a naturally-occurring, nonfeminizing, diastereomer of 17β-E2, for the treatment of liver diseases.

*Added value of this study:* In this study, using tracer-based labeling approaches in male mice subjected to CCl_4_-induced liver fibrosis, we show that 17α-E2 reduces liver fibrosis by attenuating collagen synthesis and enhancing collagen degradation mechanisms. Both transforming growth factor β1 (TGF-β1) and lysyl oxidase-like 2 (LOXL2) protein content in liver were reduced by 17α-E2. We also found that 17α-E2 increased matrix metalloproteinase 2 (MMP2) activity and suppressed stearoyl-coenzyme A desaturase 1 (SCD1) protein levels, the latter of which has been linked to the resolution of liver fibrosis. We also found that 17α-E2 increased liver fetuin-A protein, a strong inhibitor of TGF-β1 signaling, and reduced pro-inflammatory macrophage activation and cytokine expression in the liver.

*Implications of all the available evidence:* This study supports the idea that estrogens are protective against chronic liver diseases and that 17α-E2 may have therapeutic utility for the treatment of liver fibrosis.

## 1. INTRODUCTION

Nonalcoholic fatty liver disease (NAFLD) is the most common liver disease in the world^1^. Nearly 30% of the population in westernized countries have NAFLD and this number continues to increase due to rising cases of obesity and type 2 diabetes (T2D)^2^. NAFLD encompasses a spectrum of disorders ranging from simple steatosis to nonalcoholic steatohepatitis (NASH), which is characterized by lobular inflammation and hepatocyte ballooning in the presence or absence of fibrosis^3^. NASH is a major risk factor for progression to cirrhosis and hepatocellular carcinoma^4^ and represents the fastest growing indication for liver transplantation in the US over the past two decades^5^. The severity of fibrosis in the setting of NASH was recently found to be the best predictor of mortality in patients with NAFLD^6^.

Liver fibrosis manifests as an excessive accumulation of extracellular matrix (ECM) proteins, predominantly collagen, in response to chronic liver injury. Hepatic stellate cells (HSCs), a residential perisinusoidal mesenchymal cell-type, maintains ECM homeostasis in the liver. During liver injury, quiescent HSCs transdifferentiate into an activated state and develop a profibrogenic phenotype that promotes fibrosis if unabated^7^. Activated HSCs also produce transforming growth factor β1 (TGF-β1), which maintains HSC activation and is therefore the dominant mechanistic driver of liver fibrosis^7-9^. The severity of liver fibrosis is partially controlled by the degree of collagen crosslinking because it increases resistance to proteolytic degradation. Collagen crosslinking is regulated by the activity of lysyl oxidase-like 2 (LOXL2)^10,11^ in liver. The degradation of liver collagen by matrix metalloproteinases (MMPs) also plays a critical role in the control of collagen accumulation. Both MMP2 and MMP9 degrade collagen in the liver, although it remains debated if either can be targeted to enhance collagen degradation^12-17^.

Notably, the prevalence and severity of NAFLD, NASH, and liver fibrosis are greater in men than women^18-20^ and men are two-fold more likely to die from chronic liver disease^21^. However, the prevalence and severity of NAFLD, NASH, and liver fibrosis becomes comparable between the sexes following menopause, particularly when hormone replacement therapies (HRT) are not initiated^22^. Moreover, the duration of estrogen deficiency in postmenopausal women is associated with greater severity of liver fibrosis^23^. These studies suggest that estrogen signaling is protective against liver disease onset and progression. Studies in rodents support the idea that estrogens, particularly 17β-estradiol (17β-E2), are protective against hepatic lipid deposition and fibrogenesis^24-28^. However, chronic administration of 17β-E2 or combination HRTs are unrealistic in men due to increased stroke risk^29^, prostate cancer development^30^, and feminization^31^. A subset of the female population also develops side effects with chronic HRT administration including headaches, nausea, and increased risk for breast cancer and cardiovascular events^32,33^. These observations, coupled with the fact that no medications are currently approved in the US or Europe for the treatment of NAFLD, NASH, or liver fibrosis^34,35^, have led to the investigation of nonfeminizing estrogens for the treatment of liver diseases. Both naturally-occurring and synthetic nonfeminizing estrogens have been studied in recent years for their ability to elicit health benefits^36,37^.

17α-estradiol (17α-E2), a naturally-occurring diastereomer of 17β-E2 with considerably less binding affinity for classical estrogen receptors (ERα & ERβ), was recently shown to extend lifespan of male mice in a dose-dependent manner^38,39^. We and others have since reported that 17α-E2 treatment reduces adiposity and improves a myriad of systemic metabolic parameters in obese and old male mice without inducing overt feminization^40-45^. In liver, 17α-E2 dramatically reduces steatosis and age-related DNA damage, while also significantly improving hepatic insulin sensitivity in an ERα-dependent manner^40,46^. We also found that 17α-E2 attenuates several histological and transcriptional markers of liver fibrosis in models of diet-induced obesity^46^. Given that 17α-E2 dramatically reduces adiposity, it remains unclear if the benefits of 17α-E2 on liver fibrosis are secondary responses to the reversal of metabolic sequela associated with obesity, or if 17α-E2 has direct effects on fibrogenesis-related mechanisms.

In the current study, we evaluated the effects of 17α-E2 treatment on collagen synthesis and degradation rates using novel tracer-based labeling approaches in male mice subjected to CCl_4_-induced liver fibrosis. We also assessed the effects of 17α-E2 on markers of HSC activation, collagen crosslinking, and collagen degradation. Lastly, we also characterized the effects of 17α-E2 treatment of liver macrophage content and polarity due to their established role in exacerbating liver fibrosis^47,48^. We found that mice receiving 17α-E2 had significantly reduced collagen synthesis rates and greater collagen degradation rates, which was mirrored by TGF-β1 and LOXL2 protein content. These improvements were associated with increased MMP2 activity and suppressed stearoyl-coenyzme A desaturase 1 (SCD1) protein levels, the latter of which has been linked to fibrosis resolution^49,50^. We also found that 17α-E2 increased fetuin-A protein, a strong inhibitor of TGF-β1 signaling^51-55^, and reduced macrophage activation and proinflammatory markers in the liver. We conclude that 17α-E2 acts in a multimodal fashion to reduce fibrotic burden in the liver.

## 2. METHODS

### 2.1. Animal experiments

Ten-week-old male C57BL/6J mice were purchased from The Jackson Laboratory (Bar Harbor, ME, USA) and were acclimatized to the Oklahoma City VA Health Care System vivarium for a period of two weeks. Mice were individually housed with ISO cotton pad bedding, cardboard enrichment tubes, and nestlets at 22 ± 0.5°C on a 12:12-hour light-dark cycle. Unless otherwise noted, all mice had *ad libitum* access to food and water throughout the experimental timeframe. At twelve weeks of age, mice were randomized by body mass into one of five groups (**Fig. 1**), (1) Short-term carbon tetrachloride (CCl_4_) (n=18): chow-fed (TestDiet 58YP; 66.6% CHO, 20.4% PRO, 13.0% FAT), CCl_4_-treated twice weekly for four weeks, (2) Vehicle (n=15): chow-fed, vehicle-treated (olive oil) controls, (3) CCl_4_ (n=15): chow-fed, CCl_4_-treated twice weekly for eight weeks, (4) CCl_4_ + 17α preventive (n=18): chow+17α (14.4 mg/kg; Steraloids, Newport, RI)-fed for eight weeks while simultaneously being CCl_4_-treated twice weekly for eight weeks, or (5) CCl_4_ + 17α therapeutic (n=15): chow+17α-fed for the final four weeks while simultaneously being CCl_4_-treated twice weekly for eight weeks. CCl_4_ was administered as a 40% solution by i.p. injection (1 ul/g body mass) on the first and fourth day of each week as previously described^56-58^. We also administered deuterium oxide (D_2_O) in these studies so that collagen turnover rates could be evaluated. The Short-term CCl_4_ treatment group received 99% D_2_O i.p. injections at day 0 to quickly enrich the body water pool (assumed to be 60% of body mass) to 5%, which was followed by 8% D_2_O in the drinking water to maintain a steady state of body water enrichment through the remainder of the labeling period^59^. Mice in the Short-term CCl_4_ treatment group were euthanized at days 0, 4, 7, 14, 21, and 28 (n=3/time point) over the four-week intervention period. This labeling strategy allowed us to determine how changes in collagen synthesis rates contribute to increases in collagen concentration over the first 4 weeks of CCl_4_ administration. The Vehicle, CCl_4_, CCl_4_ + 17α preventive, and CCl_4_ + 17α therapeutic treatment groups were labeled for 28, 21, 14, 7, or 0 days (n=3/time point/group) prior to the end of the eight-week intervention. This labeling strategy allowed us to determine how CCl_4_ and 17α treatments impacted collagen synthesis and whether the treatments increased or decreased the pool of collagen that was resistant to turnover (i.e. fibrotic). At the conclusion of their respective interventional periods, mice were anesthetized with isoflurane and euthanized by exsanguination due to cardiac puncture. Blood was collected into EDTA-lined tubes, plasma was collected and frozen, and the mice were then perfused with ice-cold 1X PBS prior to tissues being excised, weighed, flash-frozen and store at -80°C unless otherwise noted. Following excision, small pieces of liver in the portal triad region were dissected and fixed in 4% PFA in preparation for paraffin- or cryo-embedding for future analyses. All animal procedures were reviewed and approved by the Institutional Animal Care and Use Committee of the Oklahoma City VA Health Care System.

**Fig. 1.**
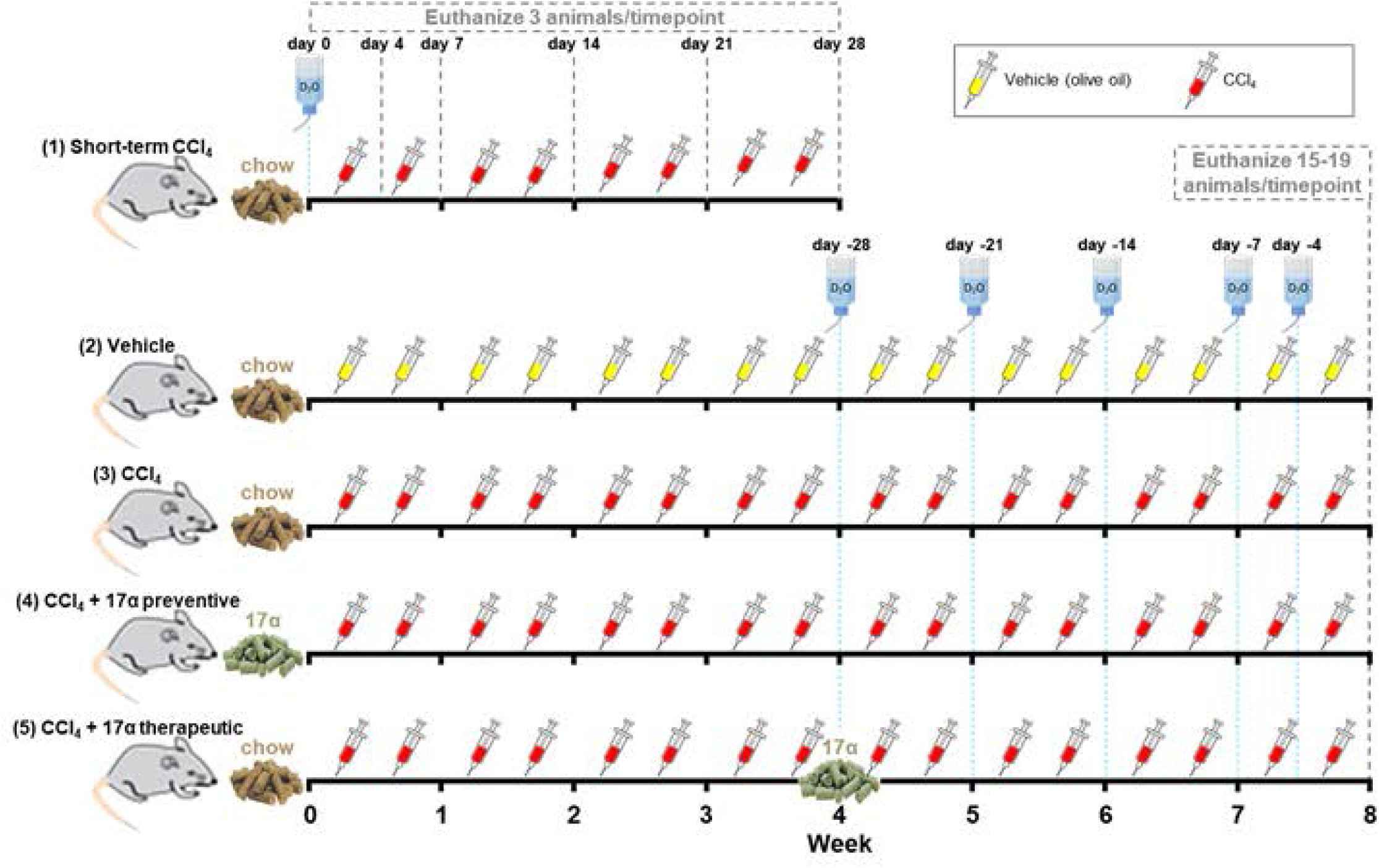
Overview of Experimental Design. Water bottles represent when 99% D_2_O bolus i.p. injections were provided and 8% D_2_O was added to the drinking water. Yellow and red syringes represent when vehicle and CCl_4_ injections were performed, respectively.

### 2.2. Plasma alanine and aspartate transaminase analyses

Plasma alanine aminotransferase (ALT) and aspartate aminotransaminase (AST) were evaluated using commercially available colorimetric kits from BioAssay Systems (Hayward, CA).

### 2.3. Liver triglyceride analyses

Liver samples (∼50 mg) were homogenized on ice for 60 seconds in 10X (v/w) RIPA Buffer (Cell Signaling, Danvers, MA) with protease and phosphatase inhibitors (Boston BioProducts, Boston, MA). Total lipids were extracted from 100 ul of homogenate using the Folch method^60^. Samples were dried under nitrogen gas at room temperature prior to being resuspended in 100μl of tert-butyl alcohol-methanol-Triton X-100 solution (3:1:1). Triglycerides (TG) were quantified spectrophotometrically using Free Glycerol & Triglyceride agents (Sigma-Aldrich, St. Louis, MO) as previously described^61^.

### 2.4. Liver histology and pathology assessments

Liver samples were fixed in 4% PFA for 24 hours, transferred to 1X PBS for 48 hours, and then transferred to 70% EtOH until paraffin embedding occurred. H&E and Masson’s trichrome staining were performed by the OMRF Imaging and Histology Core Facility using established protocols. Images of H&E and trichrome stained slides were taken on an Olympus CX43 microscope and were evaluated by two clinical pathologists who were blinded to the treatment groups. NAFLD activity scores (NAS) and fibrosis staging were determined according to NASH Clinical Research Network standards^62,63^.

### 2.5. Liver TGF-β1 quantification

Liver samples (∼100 mg) were homogenized on ice for 60 seconds in 10X (v/w) RIPA Buffer (Cell Signaling) with protease and phosphatase inhibitors (Boston BioProducts). The homogenates were spun at 17,000 rpm for 15 minutes at 4°C and the supernatant was collected. Total protein was then quantified using a Pierce BCA kit (ThermoFisher Scientific, Waltham, MA). Supernatant (20 ul) was then diluted with TGF-β1 ELISA (Abcam, Waltham, MA) assay buffer (180 ul) which was then digested with 1N HCl (20ul) for 60 minutes at room temperature. The samples were then neutralized with 1N NaOH (20 ul) and were then further diluted with ELISA assay buffer to a total volume of 1200 ul (1:60 dilution). The samples were then evaluated according manufacturer instructions. TGF-β1 concentrations were normalized to total protein and expressed as ng/mg protein.

### 2.6. Collagen extraction and isotopic labeling analyses

We performed collagen isolation and deuterium isotopic enrichment of collagen according to our previously published methods^59^. Extracted collagen was derivatized for analysis of deuterium enrichment of alanine using Gas Chromatography-Mass Spectroscopy (7890A GC-Agilent 5975C MS, Agilent, Santa Clara, CA). To determine the precursor pool enrichment, plasma samples were prepared for analysis of deuterium enrichment on a liquid water isotope analyzer (Los Gatos Research, Los Gatos, CA, USA). The precursor enrichment of alanine was then calculated by mass isotopomer distribution analysis. The deuterium enrichments of both the protein (product) and the precursor were used to calculate fraction new: Fraction new = *E*_product_/*E*_precursor_, where the *E*_product_ is the enrichment (*E*) of protein-bound alanine and *E*_precursor_ is the calculated maximum alanine enrichment from equilibration of the body water pool. The fraction new data were then plotted across the time points and curves were fit to the data using one-phase associations. Two parameters of interest were then calculated from the curves using Graphpad Prism 9 and the one-phase association function. The software calculates rate parameter (*k*, 1/day), which reflects the protein synthetic rate, and plateau fraction new (*p*), which represents the proportion of the protein pool that is actively turning over (i.e. the dynamic protein pool), with 1.0 equal to 100% of the protein pool. Finally, rates of synthesis were multiplied by the concentration of collagen (described below) to calculate absolute rates of protein turnover.

### 2.7. Hydroxyproline analyses

Powdered liver samples (∼200 mg) were evaluated for hydroxyproline content as previously described^64^. In brief, samples were hydrolyzed in 500 μl of 6 M HCl at 105°C overnight. Samples were then vigorously vortexed and 10 ul of hydrolysate was mixed with 150 μl of isopropanol and 75 μl of acetate citrate buffer containing 1.4% chloramine-T, were vigorously vortexed, and then left to oxide for 10 minutes at room temperature Samples were then mixed with 1 ml of a 3:13 solution of Ehrlich reagent (1.5 g of 4-(dimethylamino)benzaldehyde, 5 ml ethanol, 337 μl sulfuric acid) to isopropanol and incubated for 45 minutes at 55 °C. Quantification was determined by extinction measurement of the resulting solution at 558 nm. A standard curve (0–1,000 μM, trans-4-hydroxy-L-proline) was included in each assay. Results are reported as μg hydroxyproline/mg tissue.

### 2.8. Quantitative real-time PCR

Total RNA was extracted using Trizol (Life Technologies, Carlsbad, CA) and RNA (2μg) was reverse transcribed to cDNA with the High-Capacity cDNA Reverse Transcription kit (Applied Biosystems, Foster City, CA). Real-time PCR was performed in a QuantStudio 12K Flex Real Time PCR System (ThermoFisher Scientific, Waltham, MA) using TaqMan™ Gene Expression Master Mix (ThermoFisher Scientific) and predesigned gene expression assays with FAM probes from Integrated DNA Technologies (Skokie, Illinois). Target gene expression was expressed as 2^−ΔΔCT^ by the comparative CT method^65^ and normalized to the expression of TATA-box binding protein (TBP).

### 2.9. Western blot analyses

Liver samples (∼100 mg) were homogenized on ice for 60 seconds in 10X (v/w) RIPA Buffer (Cell Signaling) with protease and phosphatase inhibitors (Boston BioProducts). The homogenates were spun at 17,000 rpm for 15 minutes at 4°C and the supernatant was collected. Total protein was then quantified using a Pierce BCA kit (ThermoFisher Scientific). Proteins were separated on Any kD Criterion TGX Stain-Free gels (Bio-Rad, Hercules, CA) at 75V for 150 minutes in running buffer (Cell Signaling). Protein was then transferred to 0.2 μm pore-size nitrocellulose membranes (Bio-Rad) at 75V for 90 minutes on ice. Primary antibodies used were MMP-9 (Novus Biologicals, Centennial, CO; 1:1000), MMP-2 (Cell Signaling; 1:1000), TIMP-1 (Sino Biological, Houston, TX; 1:2000), LOXL2 (Novus Biologicals; 1:2000), PPARγ (Cell Signaling; 1:1000), SCD-1 (Cell Signaling; 1:1000), Fetuin-A (Abcam, 1:2000), p65 (Cell Signaling; 1:1000), phospho-p65 (serine^536^; Cell Signaling; 1:1000), GAPDH (Abcam; 1:2500). Primary antibody detection was performed with IRDye 800CW Infrared Rabbit antibody (LI-COR Biotechnology, Lincoln, NE) at a concentration of 1:15000. GAPDH was diluted in 5% dry milk (Cell Signaling) and all other antibodies were diluted in 5% BSA (Cell Signaling). Imaging was done on an Odyssey Fc Imaging System (LI-COR Biotechnology) with a two-minute exposure time at 800λ. Protein quantification was performed using Image Studio Software (LI-COR Biotechnology).

### 2.10. Gelatin zymography analyses

Activity of MMP-9 and MMP-2 were determined by gelatin zymography as described previously^17^ with slight modifications. In brief, resolving gels were made with 4.6 ml sterile distilled water, 2.7 ml 30% acrylamide, 2.5 ml 1.5M Tris (pH 8.8), 100 μl 10% SDS, 285 μl 8mg/ml bovine skin type B gelatin, 6 μl TEMED, and 100 μl 10% APS to reach 8% acrylamide concentrations. Stacking gels were made with 3.4 ml sterile distilled water, 830 μl 30% acrylamide, 630 μl 1M Tris (pH 6.8), 50 μl 10% SDS, 5 μl TEMED, and 50 μl 10% APS. Liver samples (∼100 mg) were homogenized on ice for 60 seconds in 10X (v/w) RIPA Buffer (Cell Signaling) with protease and phosphatase inhibitors (Boston BioProducts). The homogenates were spun at 17,000 rpm for 15 minutes at 4°C and the supernatant was collected. Total protein was then quantified using a Pierce BCA kit (ThermoFisher Scientific, Waltham, MA). Samples were prepared using non-reducing sample buffer and were loaded (20 ug/well) onto the gelatin gel along with recombinant MMP9 (Abcam) and MMP2 (BioLegend, San Diego, CA) as a standard control and run at 75V for 150min in running buffer (Cell Signaling). The gel was carefully removed and washed with Novex renaturing buffer (ThermoFisher Scientific) for 30 minutes at room temperature and was then washed with Novex developing buffer (ThermoFisher Scientific) for 30 minutes. Gels were then placed in Novex developing buffer and incubated for 16 hours at 37°C. After incubation, gels were stained with Coomassie Brilliant Blue solution (0.5 g Brilliant Blue, 250 ml methanol, 100 ml acetic acid, 150 ml sterile distilled water) for 1 hour, then washed with destaining solution (400 ml methanol, 100 ml acetic acid, 500 ml sterile distilled water) until transparent bands were visualized in the blue background. Gels were then scanned and band intensity representing MMP activity was calculated using ImageJ densitometry software.

### 2.11. Immunofluorescence analyses

Cryo-embedded liver samples were sectioned (10 μm) and stained with primary antibodies against F4/80 (total macrophages; USBiological Life Sciences; 1:250), CD11c (M1: pro-inflammatory macrophages; Invitrogen; 1:300), and CD206 (M2: anti-inflammatory macrophages; Cell Signaling; 1:500) as previously described^66^. Secondary antibodies used include goat anti-Armenian hamster IgG Alexa Fluor 594 (Jackson ImmunoResearch Laboratories, West Grove, PA; 1:500), goat anti-chicken IgG Alexa Fluor 647 (Jackson ImmunoResearch Laboratories; 1:500), and goat anti-rabbit IgG Alexa Fluor 488 (Jackson ImmunoResearch Laboratories; 1:500). Sections were mounted in Prolong Diamond Mounting Medium with DAPI (Abcam) and images were acquired using a Leica 3D Thunder scope from three non-intersecting fields per mouse. Intensity of fluorescence was measured as percent of total area using Image J after each image had its background intensity subtracted out

### 2.12. Statistical analyses

Results are presented as mean ± SEM with *p* values less than 0.05 considered significantly different. Analyses of differences between groups were performed by paired Student’s t-test or one-way ANOVA with Tukey post-hoc comparisons where appropriate using GraphPad Prism Software, Version 9.

## 3. RESULTS

### 3.1. 17α-E2 treatment attenuates markers of liver injury

To investigate the effects of 17α-E2 on the development of liver fibrosis in mice, we administered CCl_4_ to induce hepatotoxic stress while also providing 17α-E2 for the entire duration of the eight-week study (preventive group) or the final four weeks of the study (therapeutic group). Consistent with our previous reports^40-43,46,67,68^, 17α-E2 treatment reduced body mass at the conclusion of the study in both the preventive and therapeutic groups (**Fig. 2a**). As expected, CCl_4_ administration resulted in liver damage as evidenced by increased circulating ALT and AST, which were significantly blunted in both the preventive and therapeutic groups following 17α-E2 treatment (**Fig. 2b-c**). Liver mass and TG content were mildly increased by CCl_4_ administration (**Fig. 2d-e**), which was prevented by eight weeks of 17α-E2 treatment but not four weeks of therapeutic treatment (**Fig. 2e**). Pathological liver assessment confirmed liver damage, with CCl_4_ administration dramatically increasing the NASH Activity Score (NAS) and Brunt Fibrosis Score as compared to vehicle treated controls (**Fig. 2f-g**). Interestingly, 17α-E2 treatment only improved steatosis and inflammation in the preventive group, whereas both the preventive and therapeutic groups displayed declines in the severity of fibrosis following treatment (**Fig. 2 f-g**). Based on these observations, we subsequently evaluated the effects of 17α-E2 treatment on ECM homeostasis.

**Fig. 2.**
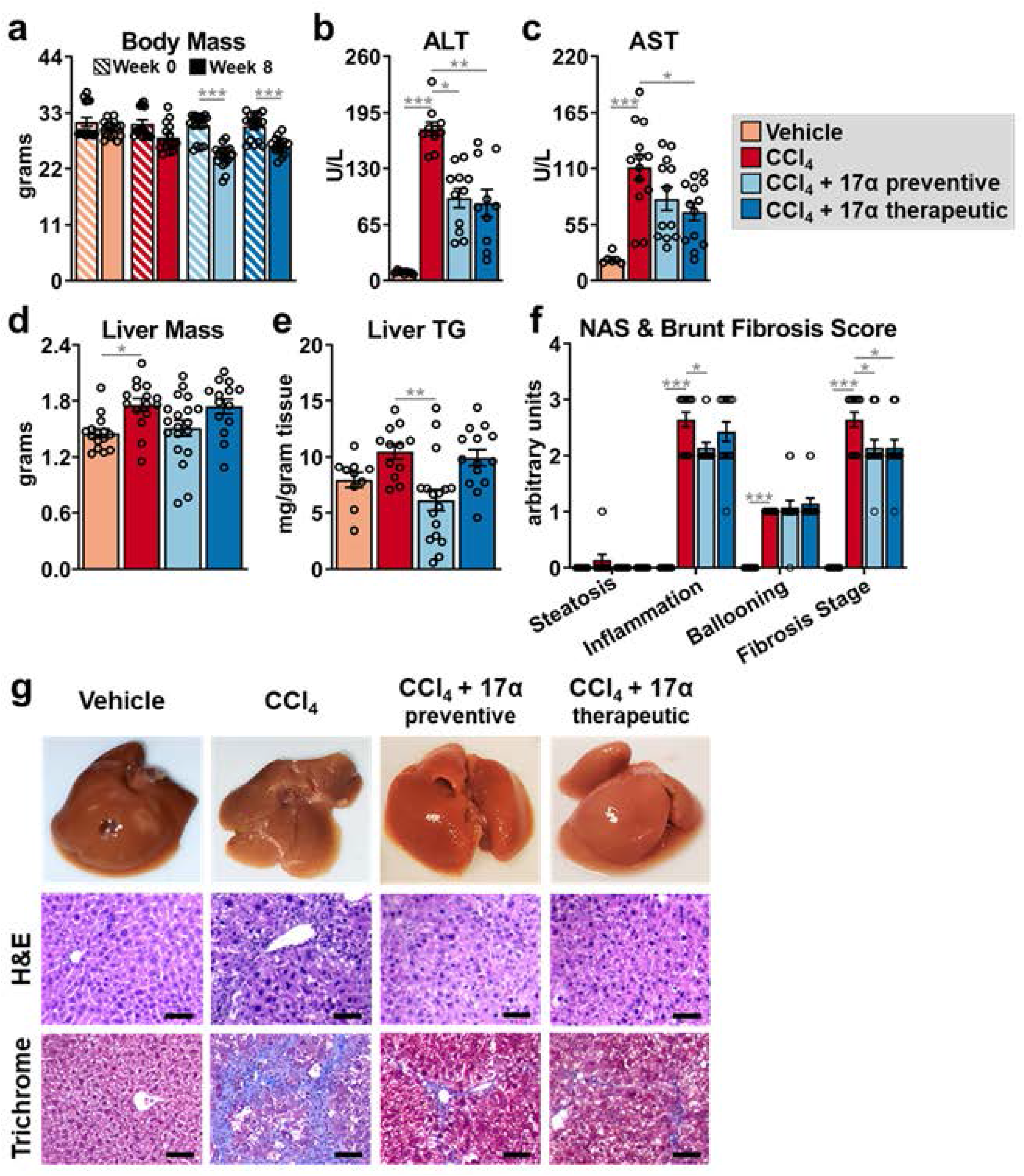
17α-E2 treatment attenuates markers of liver injury. (a) Body mass at baseline (week 0; striped) and the conclusion (solid) of the 8-week intervention [n=15-19/group]. (b) Plasma ALT [n=9-12/group], (c) plasma AST [n=6-14/group], (d) liver mass [n=15-19/group], (e) liver TG [n=11-17/group], and (f) liver pathological NASH Activity and Brunt Fibrosis Scoring at the conclusion of the 8-week intervention [n=10-14/group]. (g) Representative images of gross morphology, H&E stained (magnification = 20X; scale bar = 50 μm), and Masson’s trichrome stained (magnification = 20X; scale bar = 50 μm) liver at the conclusion of the 8-week intervention. All data are shown as mean ± SEM and were analyzed by paired Student’s t-test within treatment group (a) or one-way ANOVA with Tukey post-hoc testing (b-f). We did not indicate statistical differences between vehicle and 17α-E2 treatment groups (preventive and therapeutic), or between 17α-E2 treatment groups (preventive and therapeutic), for purposes of visual clarity. *p<0.05, **p< 0.01, ***p<0.005.

### 3.2. 17α-E2 treatment suppresses collagen synthesis in the liver

Similar to previous reports^69-72^, CCl_4_ administration significantly increased hepatic TGF-β1 production (**Fig. 3a**) and suppressed cytochrome P450 2e1 (Cyp2e1) expression (**Suppl. Fig. 1**). Cyp2e1 mediates the hepatotoxicity of CCl_4_ by metabolizing it to the free radical trichloromethyl^69,73^. These observations indicate that HSCs had become activated and developed a profibrogenic phenotype. Interestingly, 17α-E2 treatment prevented and/or reversed TGF-β1 production in the preventive and therapeutic groups, respectively (**Fig. 3a**). This observation suggests that 17α-E2 treatment may reverse HSC activation and/or promote apoptosis, which are known to occur during the resolution of liver fibrosis^74^. Notably, Cyp2e1 expression was unchanged by 17α-E2 treatment, which suggests CCl_4_ metabolism is unchanged by 17α-E2. Using stable isotope labeling techniques, we evaluated liver collagen synthesis rates. In alignment with the increase in TGF-β1 production, CCl_4_ administration robustly increased the absolute rate of collagen synthesis by nearly 3-fold (**Fig. 3b**). The preventive and therapeutic 17α-E2 treatment groups displayed collagen synthesis rates that were half of the CCl_4_ treatment group, indicating a possible prevention of an increase or a renormalization of synthesis rates (**Fig. 3b**). Liver hydroxyproline, a marker of collagen content^75^, increased following four (**Suppl. Fig. 2**) and eight weeks of CCl_4_ administration (**Fig. 3c**). 17α-E2 treatment significantly reduced hydroxyproline levels in the therapeutic group, but only induced a downward trend in the preventive group (**Fig. 3c**). By our stable isotope labeling approach, we were able to determine how much of the collagen pool was resistant to turnover (static), which we have previously shown to be indicative of a fibrotic phenotype^59^. As anticipated, the percentage of collagen in the static pool was increased with CCl_4_ administration (**Fig. 3d**), as was the absolute amount of collagen in both the static and dynamic pools (**Fig. 3e**). Introducing 17α-E2 treatment at different phases of fibrosis development led to differential responses in collagen turnover dynamics. The preventive treatment group displayed a higher percentage of dynamic collagen compared to the CCl_4_ treatment group (**Fig. 3d**), but the overall difference in absolute collagen in both the static and dynamic pools was not found to be statistically different (**Fig. 3e**). Conversely, the therapeutic group displayed a higher percentage of collagen in the static pool compared to the CCl_4_ treatment group (**Fig. 3d**), although the absolute amount of collagen in the static pool was unchanged (**Fig. 3e**). This shorter duration of 17α-E2 treatment (4 weeks) in the therapeutic group dramatically reduced the absolute amount of collagen in the dynamic pool, which was reflective of the overall lower collagen concentration (**Fig 3e**). Collectively, these findings suggest that depending on the timing of administration, 17α-E2 treatment may slow the progression of fibrosis or resolve the collagen pool that is not crosslinked. Therefore, we next sought to determine if 17α-E2 treatment alters the expression and/or activity of LOXL2 and MMPs.

**Fig. 3.**
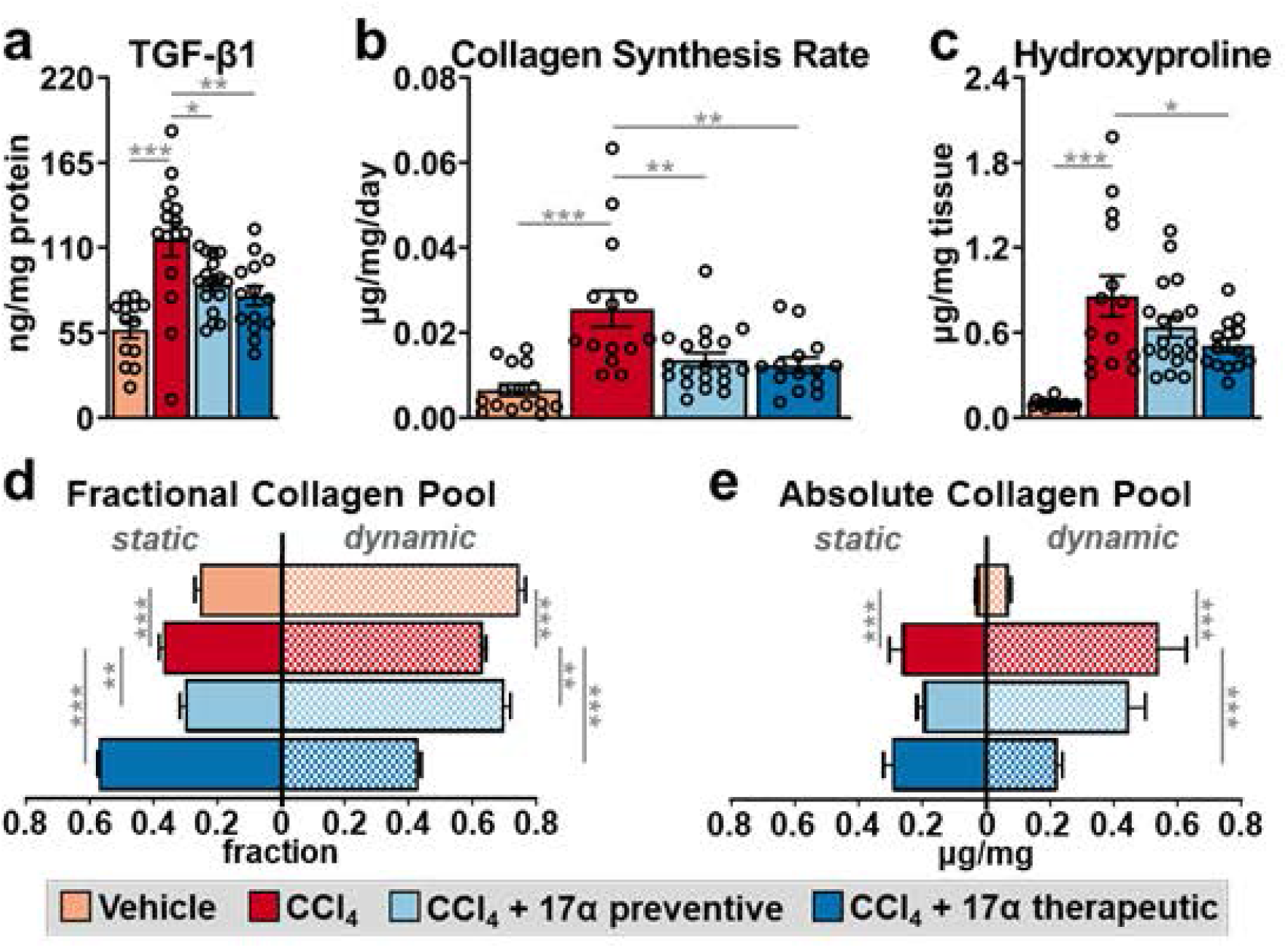
17α-E2 treatment suppresses collagen synthesis and fibrotic collagen deposition in liver. (a) Liver TGF-β1 at the conclusion of the 8-week intervention [n=13-17/group]. (b) Collagen synthesis rates during the 8-week intervention [n=14-18/group]. (c) Liver TGF-β1 hydroxyproline content at the conclusion of the 8-week intervention [n=14-19/group]. (d) Fractional percent [n=15-18/group], and (e) absolute content [n=15-18/group] of static and dynamic collagen in liver at the conclusion of the 8-week intervention. All data are shown as mean ± SEM and were analyzed by one-way ANOVA with Tukey post-hoc testing. We did not indicate statistical differences between vehicle and 17α-E2 treatment groups (preventive and therapeutic), or between 17α-E2 treatment groups (preventive and therapeutic), for purposes of visual clarity. *p<0.05, **p< 0.01, ***p<0.005.

### 3.3. 17α-E2 treatment suppresses LOXL2 expression and increased MMP2 activity

To determine if the declines in collagen synthesis following 17α-E2 treatment were due to changes in collagen crosslinking and/or degradation mechanisms, we first evaluated LOXL2 expression due to its key role in fibrotic matrix stabilization^11^. It should also be noted that previous reports have established that the magnitude of LOXL2 expression is indicative of activity, and thus, an increase in crosslinked collagen fibrils^76^. In alignment with previous reports^77,78^, CCl_4_ administration significantly increased LOXL2 expression at both the transcript and protein level (**Fig. 4a-c**), indicating a higher level of collagen crosslinking is occurring. 17α-E2 treatment did not alter *Loxl2* transcription but did mildly suppress LOXL2 protein in the therapeutic group (**Fig. 4a-c**), suggesting that a decline in collagen crosslinking may be a minor mechanism by which 17α-E2 reduces liver fibrosis. We subsequently evaluated MMP expression and activity due to their critical roles in controlling collagen degradation. We focused most of our efforts on analyzing MMP2 and MMP9 because they control the final step in collagen degradation^7^. We also evaluated the expression of tissue inhibitor matrix metalloproteinase 1 (TIMP1) due to its established role in inhibiting MMP activity^17^. We found that CCl_4_ administration increased *Mmp2* and *Timp1* transcripts, but did not alter *Mmp9* mRNA expression (**Fig. 5a-c**). CCl_4_-mediated liver fibrosis was also associated with increased transcription of *Mmp8, Mmp12, Mmp13, Mmp14, Mmp16*, and *Mmp19* (**Suppl. Fig. 3**). 17α-E2 treatment did not alter transcription of any MMP we analyzed or Timp1 (**Fig. 5a-c; Suppl. Fig. 3**). To our surprise, CCl_4_ administration did not alter MMP2 or MMP9 protein expression, but did promote declines in TIMP1 protein (**Fig. 5d-g**). 17α-E2 treatment mildly increased MMP2 protein expression, preferentially in the therapeutic group, but failed to modulate MMP9 or TIMP1 protein levels (**Fig. 5d-g**). We next sought to determine if MMP2 or MMP9 activity was altered. As expected^12,79-81^, CCl_4_ administration dramatically increased MMP2 and MMP9 activity as determined by gelatin zymography (**Fig. 5h-j**). Interestingly, 17α-E2 treatment further increased MMP2 activity in both the preventive and therapeutic groups (**Fig. 5h-j**), indicating that declines in liver collagen by 17α-E2 are at least partially mediated through degradation mechanisms.

**Fig. 4.**
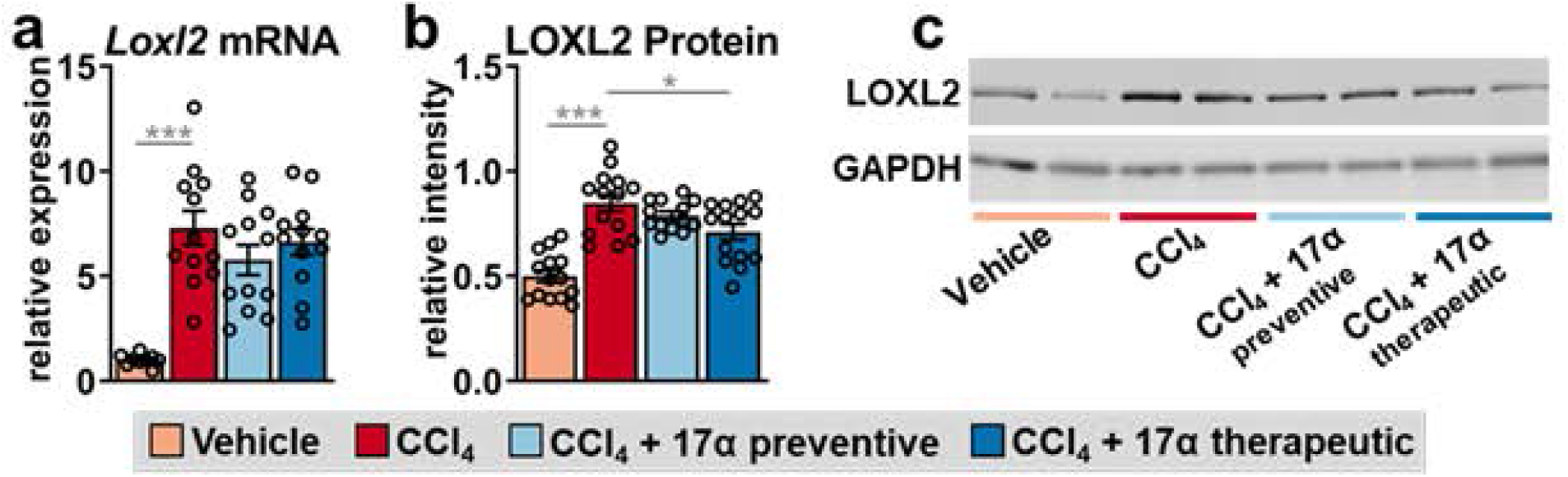
17α-E2 treatment reduces the collagen crosslinking enzyme LOXL2 in liver. Liver (a) *Loxl2* mRNA [n=9-12/group], and (b) LOXL2 protein [n=14/group] at the conclusion of the 8-week intervention. (c) Representative immunoblots of liver LOXL2 and GAPDH at the conclusion of the 8-week intervention. All data are shown as mean ± SEM and were analyzed by one-way ANOVA with Tukey post-hoc testing. We did not indicate statistical differences between vehicle and 17α-E2 treatment groups (preventive and therapeutic), or between 17α-E2 treatment groups (preventive and therapeutic), for purposes of visual clarity. *p<0.05, ***p<0.005.

**Fig. 5.**
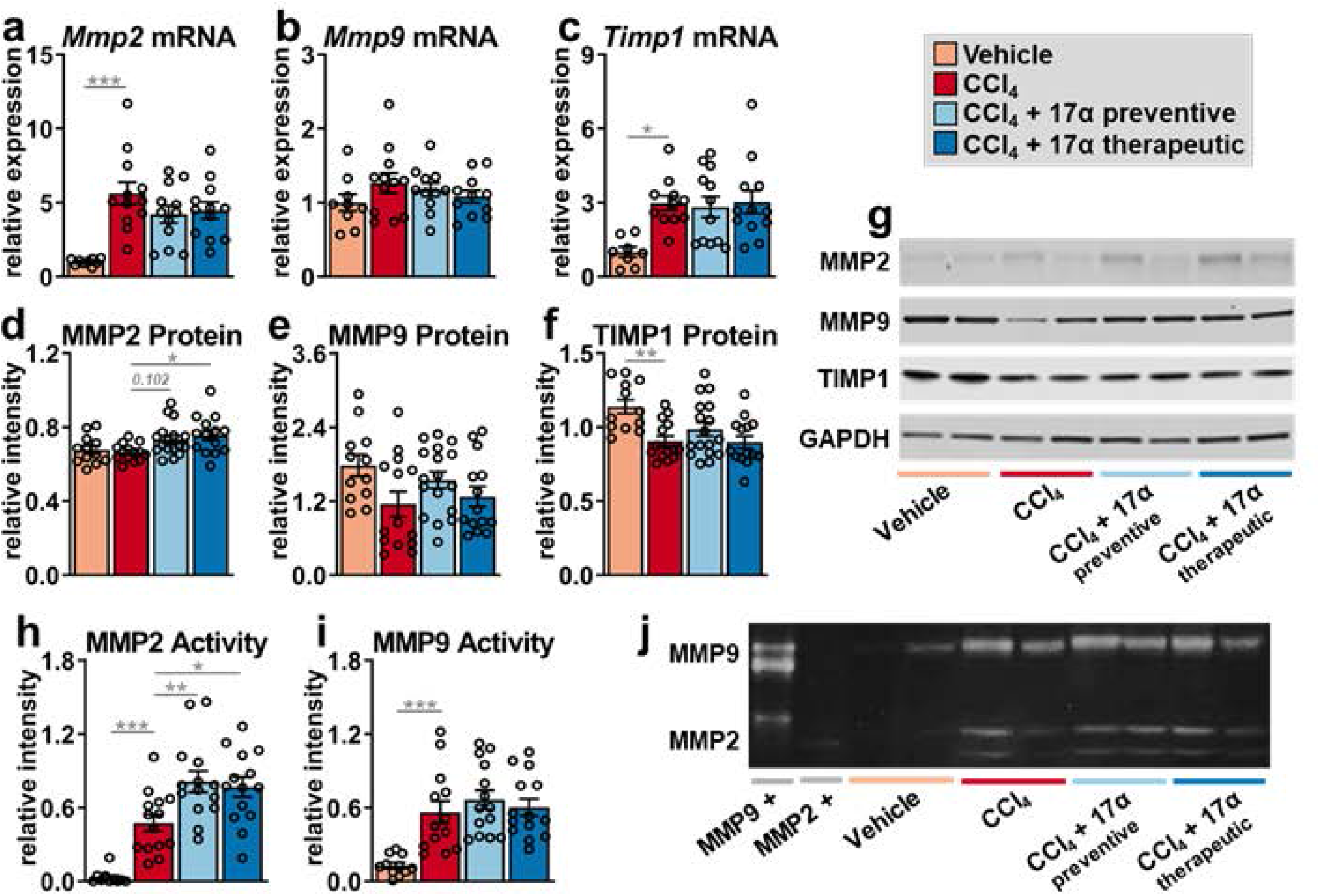
17α-E2 treatment increases collagen degradation by modulating MMP2 activity in liver. Liver (a) *Mmp2* mRNA [n=9-12/group], (b) *Mmp9* mRNA [n=9-12/group], (c) *Timp1* mRNA [n=8-11/group], (d) MMP2 protein [n=12-17/group], (e) MMP9 protein [n=12-17/group], and (f) TIMP1 protein [n=12-17/group] at the conclusion of the 8-week intervention. (g) Representative immunoblots of liver MMP2, MMP9, TIMP1, and GAPDH at the conclusion of the 8-week intervention. Liver (h) MMP2 [n=10-14/group], and (i) MMP9 [n=11-14/group] enzymatic activity at the conclusion of the 8-week intervention. (j) Representative images of liver gelatin zymography assessing MMP2 and MMP9 enzymatic activity. Signal intensity for each sample was normalized to the intensity of their corresponding positive control standard. All data are shown as mean ± SEM and were analyzed by one-way ANOVA with Tukey post-hoc testing. We did not indicate statistical differences between vehicle and 17α-E2 treatment groups (preventive and therapeutic), or between 17α-E2 treatment groups (preventive and therapeutic), for purposes of visual clarity. *p<0.05, **p< 0.01, ***p<0.005.

### 3.4. 17α-E2 treatment suppresses hepatic SCD1 expression and increases Fetuin-A expression

To determine if the declines in collagen synthesis following 17α-E2 treatment were due to changes in mechanisms linked to the resolution of liver fibrosis, we evaluated peroxisome proliferator-activated receptor-gamma (PPARγ), SCD1, and fetuin-A expression. Several reports have established that PPARγ actively inhibits HSC activation and that its expression and activity are reduced once HSC activation occurs^82-86^. Moreover, mice with HSC-specific PPARγ disruption display defects in the resolution of liver fibrosis following the discontinuation of hepatotoxic treatments^87^. We found that CCl_4_ administration did not alter *Pparγ* transcription or translation in whole liver, whereas 17α-E2 treatment increased *Pparγ* mRNA and protein in the preventive group only (**Fig. 6a, c**). In contrast to PPARγ, CCl_4_ administration significantly increased *Scd1* transcripts and protein (**Fig 6b, d**). SCD1 is the rate-limiting enzyme that catalyzes the formation of monounsaturated fatty acids and its association with liver steatosis and fibrosis has been known for years^50,88-91^. 17α-E2 treatment significantly reduced *Scd1* transcripts in the preventive group (**Fig. 6b**), which was mirrored by a robust downregulation of SCD1 protein in the preventive group (**Fig. 6d**). Interestingly, the therapeutic 17α-E2 treatment group also displayed a trending reduction in SCD1 protein level (**Fig. 6d**), suggesting that a longer treatment duration may prove beneficial when intervening in the context of established liver fibrosis. We also evaluated fetuin-A expression due to it being a strong inhibitor of TGF-β1 signaling^51-55^. We found that CCl_4_ administration did not alter fetuin-A protein levels in the liver but that both the preventive and therapeutic 17α-E2 treatment groups displayed increased levels of fetuin-A (**Fig. 7a-b**). Collectively, these observations suggest that 17α-E2 treatment modulates several pathways linked to the prevention and/or resolution of liver fibrosis.

**Fig. 6.**
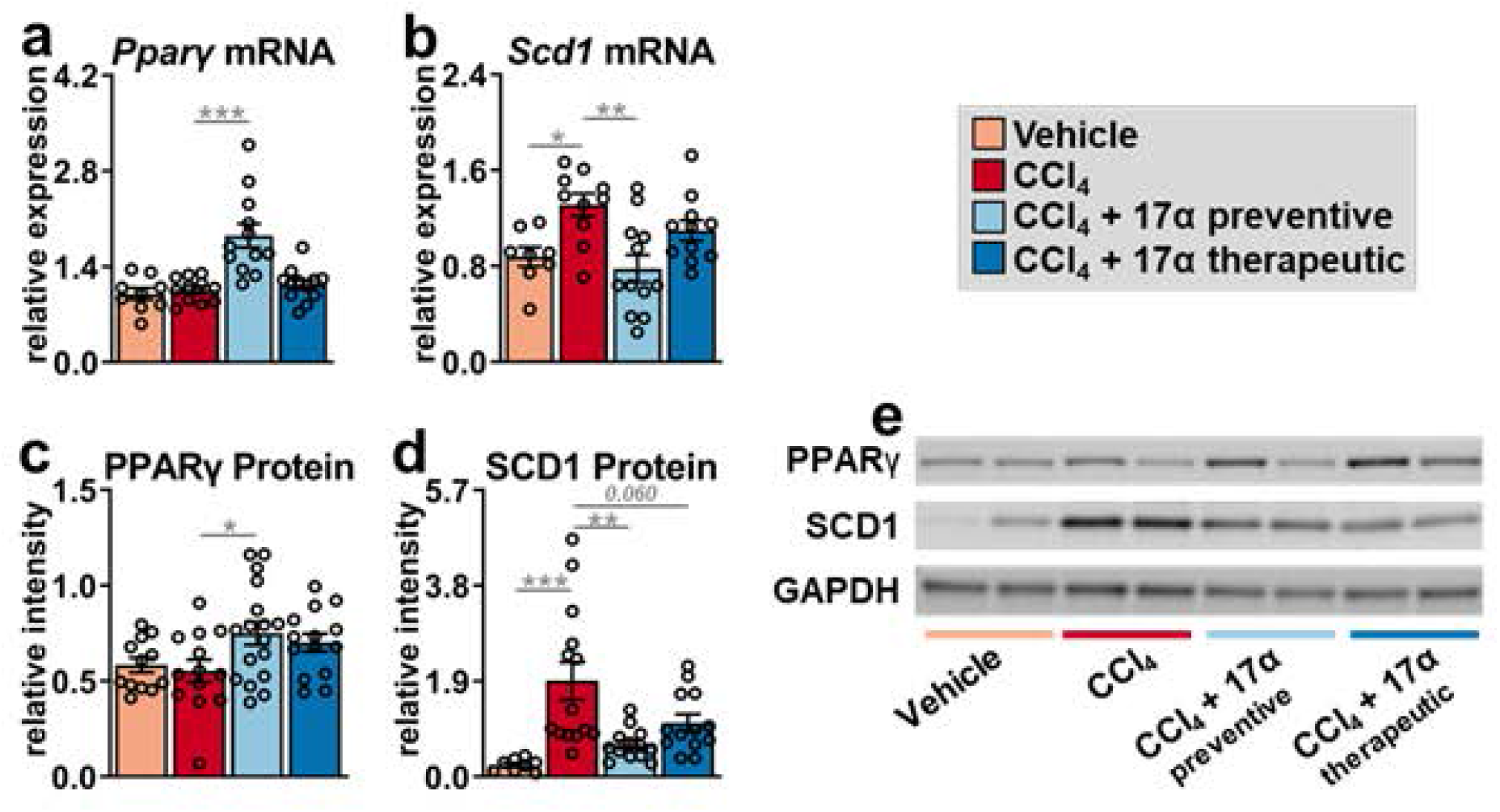
17α-E2 treatment beneficially modulates fibrosis resolution genes, PPARγ and SCD1, in liver. Liver (a) *Pparγ* mRNA [n=9-12/group], (b) *Scd1* mRNA [n=9-12/group], (c) PPARγ protein [n=12-17/group], and (d) SCD1 protein [n=8-14/group] at the conclusion of the 8-week intervention. (e) Representative immunoblots of liver PPARγ, SCD1, and GAPDH at the conclusion of the 8-week intervention. All data are shown as mean ± SEM and were analyzed by one-way ANOVA with Tukey post-hoc testing. We did not indicate statistical differences between vehicle and 17α-E2 treatment groups (preventive and therapeutic), or between 17α-E2 treatment groups (preventive and therapeutic), for purposes of visual clarity. *p<0.05, **p< 0.01, ***p<0.005.

**Fig. 7.**
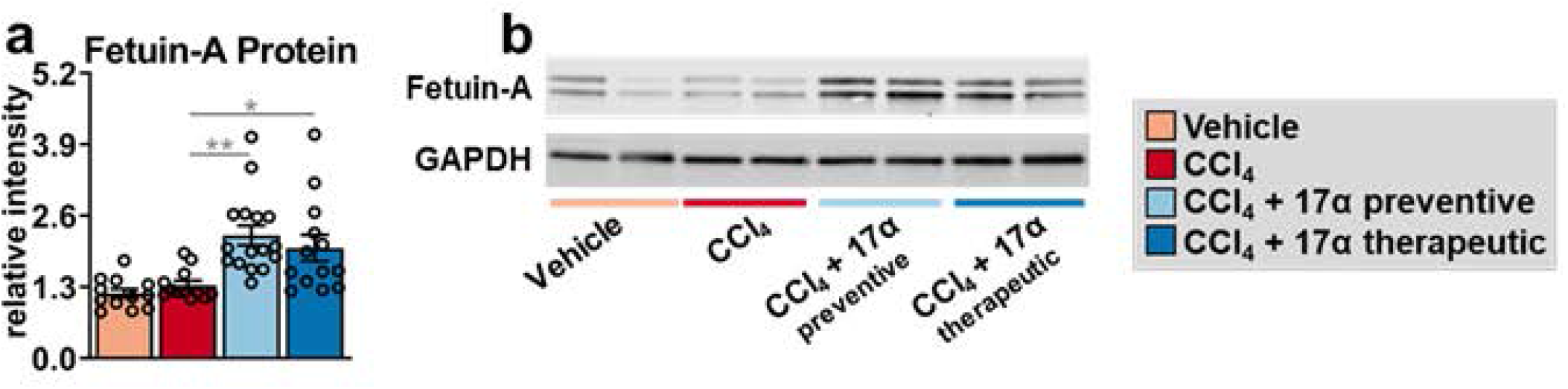
17α-E2 treatment increases fetuin-A, a potent inhibitor of TGF-β1, in liver. (a) Liver fetuin-A protein at the conclusion of the 8-week intervention [n=12-17/group]. (b) Representative immunoblots of liver fetuin-A and GAPDH at the conclusion of the 8-week intervention. All data are shown as mean ± SEM and were analyzed by one-way ANOVA with Tukey post-hoc testing. We did not indicate statistical differences between vehicle and 17α-E2 treatment groups (preventive and therapeutic), or between 17α-E2 treatment groups (preventive and therapeutic), for purposes of visual clarity. *p<0.05, **p< 0.01.

### 3.5. 17α-E2 treatment suppresses hepatic inflammation by altering macrophage content and polarity

Given the established role of macrophages in the progression and resolution of liver fibrosis^47,48^, we sought to determine if the aforementioned improvements in liver injury with 17α-E2 treatment were associated with changes in macrophage content and polarity in liver. In alignment with previous reports^66,92^, CCl_4_ administration significantly increased total macrophage (F4/80) content in the liver by over two-fold (**Fig. 8a-b**), while also increasing M1 pro-inflammatory macrophage (CD11c) content by over 10-fold (**Fig. 8a, c**). These changes were mirrored by increased transcription of *F4/80*, C-C motif chemokine ligand 5 (*Ccl5*), and C-X-C motif chemokine ligand 2 (*Cxcl2*) (**Suppl. Fig. 4a-c**); the latter two of which are chemokines linked to liver fibrosis severity^93,94^. Liver expression of cytokines, tumor necrosis factor-alpha (*Tnfα*) and interleukin 1 beta *(Il1β)*, were upregulated by CCl_4_ administration but only *Tnfα* was found to be significantly increased (**Suppl. Fig. 4d-e**). These observations suggest that both resident and recruited macrophages were activated in response to CCl_4_ administration, which has been established previously^47,48^. 17α-E2 treatment reduced total macrophage and M1 pro-inflammatory macrophage content in the liver in both treatment groups (**Fig. 8a-c**), although these reductions were only found to be significantly different in the preventive treatment group; suggesting that 17α-E2 treatment duration likely plays a major role in the suppression of pro-inflammatory macrophage activity. Interestingly, CCl_4_ administration also mildly increased M2 anti-inflammatory macrophage content in the liver (**Fig. 8a, d**), which has previously been described as a compensatory response to severe liver injury^92,95^. 17α-E2 treatment further increased M2 anti-inflammatory macrophage content in the liver by nearly two-fold in both the preventive and therapeutic groups (**Fig. 8a, d**), indicating that 17α-E2 robustly alters liver macrophage polarity within a few weeks of treatment. 17α-E2 treatment also suppressed *F4/80, Ccl5, Cxcl2, Tnfα, and Il1β* mRNA to varying degrees in both the preventive and therapeutic groups (**Suppl. Fig. 4a-e**). These observations suggest that the resolution of liver fibrosis with 17α-E2 treatment may be at least partially mediated through actions in immune cells. For additional confirmation that 17α-E2 reduced the liver pro-inflammatory milieu, we evaluated the activation of the NF-κB subunit p65, which is commonly upregulated in chronic liver injury and fibrosis^96,97^. We found that CCl_4_ administration increased liver p65 serine^536^ phosphorylation by over three-fold (**Fig. 8e-f**), which represents the phosphorylation site with the most potent inducible response to inflammatory stimuli^98^. 17α-E2 completely reversed this induction in both the preventive and therapeutic treatment groups (**Fig. 8e-f**), indicating that 17α-E2 can reverse liver inflammation within a few weeks of treatment. Collectively, these observations suggest that 17α-E2 treatment may attenuate liver fibrosis by suppressing pro-inflammatory responses in the liver.

**Fig. 8.**
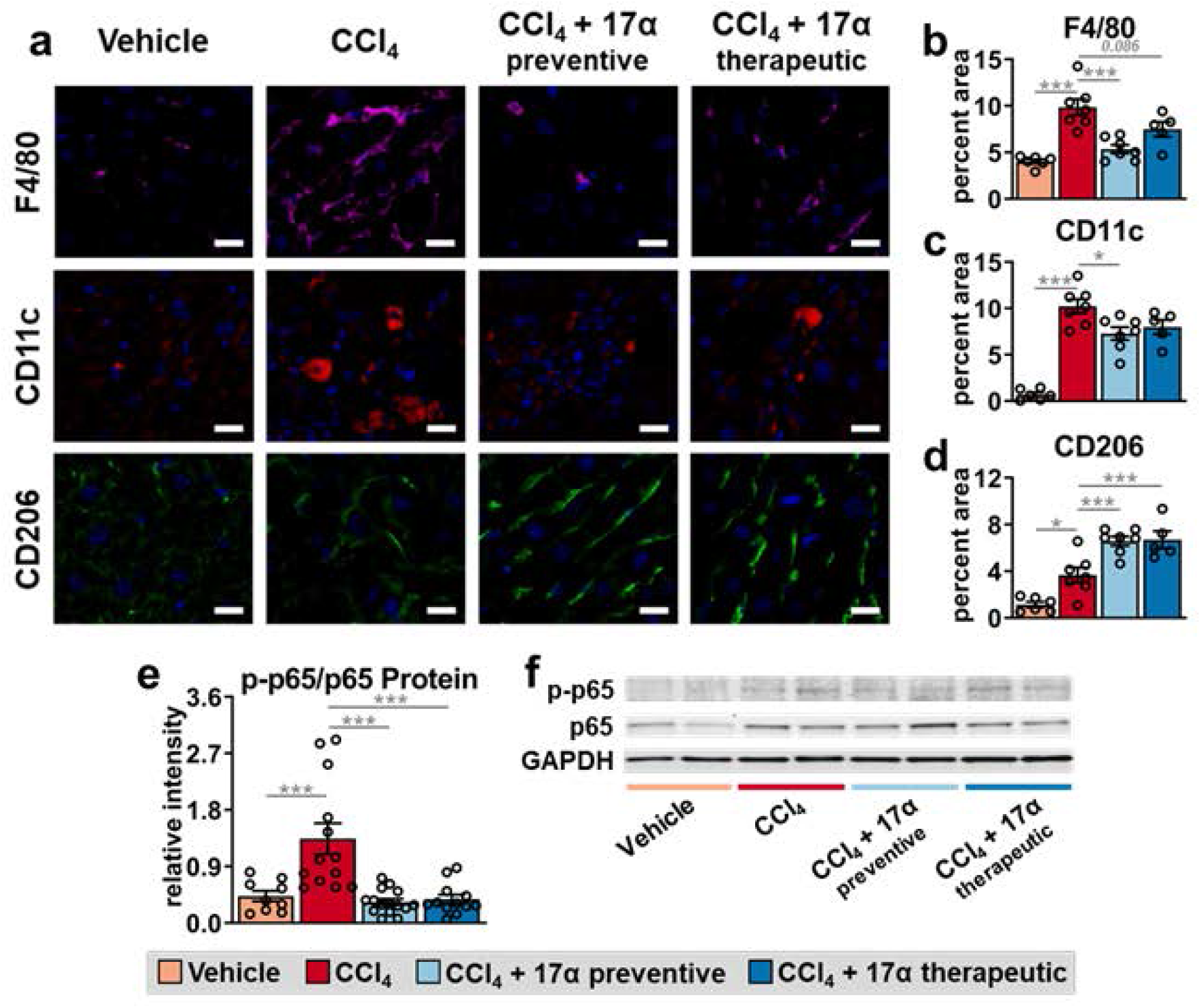
17α-E2 treatment attenuates macrophage infiltration and proinflammatory responses in liver. (a) Representative immunofluorescence images of F4/80 (total macrophages), CD11c (M1: pro-inflammatory macrophages), and CD206 (M2: anti-inflammatory macrophages) in liver at the conclusion of the 8-week intervention (magnification = 320X; scale bar = 50 μm). Percentage of liver area for (b) F4/80 [n=5-7/group], (c) CD11c [n=5-7/group], and (d) CD206 [n=5-7/group] at the conclusion of the 8-week intervention. (e) Ratio of liver phospho to total p65 at the conclusion of the 8-week intervention [n=12-16/group]. (f) Representative immunoblots of liver phosphor p65, total p65, and GAPDH at the conclusion of the 8-week intervention. All data are shown as mean ± SEM were analyzed by one-way ANOVA with Tukey post-hoc testing. We did not indicate statistical differences between vehicle and 17α-E2 treatment groups (preventive and therapeutic), or between 17α-E2 treatment groups (preventive and therapeutic), for purposes of visual clarity. *p<0.05, ***p<0.005.

## 4. DISCUSSION

17α-E2 is a naturally-occurring diastereomer of 17β-E2 that ameliorates metabolic dysfunction in obese and aged male mice^40-42,46,68^, which we surmise underlies its lifespan-extending effects^38,39^. We have previously reported that 17α-E2 reverses hepatic steatosis and improves hepatic insulin sensitivity in an ERα-dependent manner^46^. Given that hepatic steatosis and insulin resistance are directly linked to the onset of liver fibrosis^16,35^, this study aimed to determine if 17α-E2 treatment could beneficially modulate fibrogenesis-related mechanisms in the absence of obesity and hepatic steatosis. Herein, we clearly demonstrate that preventive and therapeutic 17α-E2 treatments attenuate or reverse the progression of liver fibrosis by modulating collagen turnover. Collectively, our data suggests that 17α-E2 likely acts in a multimodal fashion through several cell-types to attenuate liver injury and fibrosis.

One of the most important findings from this study was the observation that 17α-E2 significantly suppressed collagen synthesis in the setting of chronic liver injury without hepatic steatosis. This indicates that 17α-E2 reduces fibrogenesis through mechanisms other than the reduction of lipotoxicity. One such potential mechanism is the prevention and/or reversal of HSC activation, which is supported by our data showing declines in liver TGF-β1 production and increased liver fetuin-A production following 17α-E2 treatment. Fetuin-A antagonizes TGF-β1 binding to its receptor and modulates downstream Smad pathway activity, thereby inhibiting HSC activation^52-54^. Potential HSC inactivation by 17α-E2 directly, or through fetuin-A mediated mechanisms, is further supported by the data generated through labeling of liver collagen, which revealed that 17α-E2 treatment predominantly reduced the dynamic pool of collagen in the liver. This finding suggests that collagen production by HSCs was attenuated and that the degree of collagen crosslinking of LOXL2 was also reduced. This was particularly evident in the therapeutic treatment group that received 17α-E2 for the final four weeks of the intervention period. Interestingly, the therapeutic treatment group did not show declines in the static pool of liver collagen, indicating that four weeks of 17α-E2 treatment is insufficient to induce the unwinding and degradation of stabile collagen fibrils. This is not incredibly surprising given the resistance of stabile collagen fibrils to proteolytic degradation^99^, coupled with the fact that hepatic injury due to CCl_4_ administration was ongoing during 17α-E2 treatment. Conversely, the preventive treatment group, which received 17α-E2 for the entire eight-week intervention period, displayed trending reductions in both the dynamic and static collagen pools. It is unclear if this observation indicates that concomitant 17α-E2 treatment reduced the level of stabile collagen fibril accumulation, or increased unwinding and degradation of crosslinked collagen fibrils. We speculate that the longer duration of 17α-E2 treatment in the preventive group likely did both as evidenced by reduced collagen synthesis rates and increased MMP2 activity. Future studies will be needed to determine if 17α-E2 reduces liver collagen production by directly inducing apoptosis or inactivation of HSCs, through fetuin-A related mechanisms, or if 17α-E2 attenuates hepatocyte stress and/or immune cell activation and pro-inflammatory responses which in turn reduce HSC activation.

Another interesting outcome from our study was the observation that 17α-E2 increased liver PPARγ in the absence of hepatic steatosis. The change in PPARγ, which was more robust in the preventive treatment group, has implications for the inactivation of HSCs and suppression of pro-inflammatory activity in monocytes and macrophages. Previous studies have established that PPARγ is highly expressed in quiescent HSCs and that its activity promotes apoptosis of activated HSCs and differentiation toward a quiescent phenotype^84,100^. PPARγ actions in monocytes and macrophages are similarly anti-fibrotic. PPARγ activation in M1 pro-inflammatory macrophages is anti-inflammatory by repressing NF-κB activity and the expression of downstream response genes^101^. PPARγ activation in monocytes promotes their differentiation into M2 anti-inflammatory macrophages^102^. Moreover, macrophage-specific deletion of PPARγ increases liver fibrosis and inflammation in mice subjected to CCl_4_ administration^83^. Given that PPARγ activity in hepatocytes promotes steatosis^103^, coupled with the fact that 17α-E2 reduces hepatic lipid deposition, we surmise that one of the mechanisms by which 17α-E2 attenuates liver fibrosis is by increasing PPARγ activity in HSCs and/or macrophages. It remains unclear if 17α-E2 directly modulates PPARγ activity in HSCs or monocytes/macrophages but crosstalk between estrogen receptors (ERα & ERβ) and PPARγ has been documented previously^104-106^, although most studies suggest antagonist interactions. We have previously reported that 17α-E2 can elicit genomic activity of ERβ^40^, which is dominant to HSCs^107,108^, and ERα^40,46^, which is dominant to monocytes/macrophages^109,110^. However, it remains unresolved if the actions of 17α-E2 through ERα, and potentially ERβ, are through genomic or nongenomic mechanisms. Future studies will be needed to determine if PPARγ modulation by 17α-E2 is direct and if this occurs in HSCs, monocytes/macrophages, or both.

Our study also revealed that 17α-E2 treatment reduced liver SCD1 in the absence of hepatic steatosis. The suppression of SCD1 in hepatocytes is known to reduce steatosis^88,90,91^ and 17β-E2 is reported to be a negative regulator of SCD1 expression^111-113^. This suggests that declines in hepatic lipid accumulation with 17α-E2 treatment, which we also observed in the current study, is at least partially mediated through suppression of SCD1 in hepatocytes. What remains unclear is if 17α-E2 also directly suppresses SCD1 in HSCs, which was recently linked to an inhibition of fibrogenesis^50^. As addressed above, HSCs express ERβ and 17α-E2 can elicit genomic activity through this receptor, therefore future studies should explore the possibility that 17α-E2 attenuates liver fibrosis through direct actions in HSCs.

Lastly, our study revealed that 17α-E2 treatment elicited beneficial effects on liver macrophage content and polarity. Macrophages play a critical role in regulating the progression and resolution of liver fibrosis due to their plasticity and heterogeneity^47,48,114^. Macrophage phenotypes, or polarization, are driven by the local microenvironment and are broadly categorized as M1 pro-inflammatory or M2 anti-inflammatory^48,115^, although several subtypes of both M1 and M2 polarization states have been described^116,117^. Our study clearly demonstrates that 17α-E2 treatment reduces CCl_4_-mediated increases in total and M1 pro-inflammatory macrophages in liver, in addition to dramatically suppressing NF-κB activity and the expression of downstream response genes. Furthermore, 17α-E2 treatment increased M2 anti-inflammatory macrophages in liver, which have recently been shown to protect hepatocytes against apoptosis and necroptosis^92,95^. Interestingly, 17β-E2 is widely reported to promote macrophage polarization toward a M2 anti-inflammatory state in an ERα-dependent manner^118-120^. As alluded to above, we recently reported that 17α-E2 elicits beneficial metabolic effects through ERα^46^, therefore we surmise that 17α-E2 directly modulates macrophage polarization through ERα. However, other mechanisms may also be contributing to the change in macrophage phenotype following 17α-E2 treatment, including the potential modulation by PPARγ and fetuin-A. PPARγ^99,100^ and fetuin-A^121-123^ have been reported to suppress macrophage pro-inflammatory activity and promote M2 polarization and both were upregulated in liver by 17α-E2. At this juncture it remains unclear if 17α-E2 modulates PPARγ expression in macrophages, or if the rise in liver fetuin-A plays a mechanistic role in the shift toward M2 anti-inflammatory macrophage content following 17α-E2 treatment. Given the wealth of data indicating that ERα agonism in macrophages robustly alters the polarity state, we speculate that direct signaling of 17α-E2 through ERα in macrophages represents the dominant mechanism by which 17α-E2 alters pro-inflammatory outcomes. Future studies will be needed to determine if 17α-E2 directly modulates macrophage polarity to promote the resolution of liver fibrosis or if the changes in pro-inflammatory outcomes occur as secondary responses to 17α-E2 actions in hepatocytes and/or HSCs.

There are a few notable caveats to the current study that should be acknowledged. First, the primary limitation is that we are currently unable to determine if the benefits of 17α-E2 on liver fibrosis are largely mediated through a single cell-type, hepatocytes, HSCs, or macrophages. Although our data suggests that 17α-E2 likely elicits benefits through direct actions in all three cell-types, the current dataset does not allow us to make definitive conclusions due to the significant amount of crosstalk between the aforementioned cell-types in the setting of liver injury and fibrosis. Future studies utilizing hepatocyte-, HSC-, and macrophage-specific deletions of ER*α* or ERβ will provide tremendous insight into the effects of 17α-E2 on the prevention and/or resolution of liver fibrosis. Another minor limitation of our design and labeling approaches is that they did not allow us to determine if chronic 17α-E2 treatment increased the unwinding and degradation of static collagen, or if it prevents the crosslinking of dynamic collagen, thereby reducing static collagen accumulation. Having a greater understanding of how 17α-E2 alters collagen synthesis and degradation mechanisms will provide insight into the translatability of 17α-E2 into humans with liver fibrosis. Lastly, although CCl_4_ administration is an established model of liver fibrogenesis, it does not completely recapitulate the human NASH and liver fibrosis^35^. Future studies utilizing a combination approach of western diet and CCL_4_ administration should be undertaken to determine if 17α-E2 also elicits benefits in a disease state that more closely resembles human NASH and liver fibrosis^124^. We speculate that 17α-E2 will elicit even greater benefits in the model employing western diet and CCL_4_ administration due to our previous work establishing that 17α-E2 reduces hepatic steatosis and insulin resistance^40,46^.

In summary, the data presented herein are the first to show that 17α-E2 reduces liver fibrosis by attenuating collagen synthesis and enhancing collagen degradation mechanisms. Our data suggests that 17α-E2 acts directly on hepatocytes, HSCs, and macrophages to attenuate liver fibrosis, although additional studies will be needed to confirm this suspicion. Another potential avenue of investigation that remains unresolved is whether 17α-E2 acts through genomic or non-genomic mechanisms to elicit health benefits. Our current study provides critical insight into the how nonfeminizing estrogen compounds may have therapeutic potential for the treatment of chronic liver diseases.

## AUTHOR CONTRIBUTIONS

S.A.M., B.F.M., and M.B.S. conceived the project and designed the experiments. S.A.M., R.S., and S.N.M. performed the experiments with contributions from M.K., W.L., T.D.S., J.V.V.I., F.F.P. III, T.L., and W.M.F. S.A.M., W.M.F., and M.B.S. created figures and performed statistical analyses. S.A.M. and M.B.S. wrote the manuscript and all authors edited and approved the final manuscript.

## FUNDING & ACKNOWLEDGEMENTS

This work was supported by the National Institutes of Health (R01 AG059430 to W.M.F., R56 AG067754 to B.F.M., R01 AG070035 to M.B.S.) and the US Department of Veterans Affairs (I01 BX003906 & ISI BX004797 to W.M.F., Pilot Research Funding to M.B.S.). We thank Ms. Catelyn Jones and Ms. Kennedy Gosnell for animal colony management and Ms. Claire Abbott for assistance with sample preparation.

## SUPPLEMENTARY MATERIALS

Supplementary data associated with this article can be found online.

## FIGURE LEGENDS

**Supplemental Figure 1:**
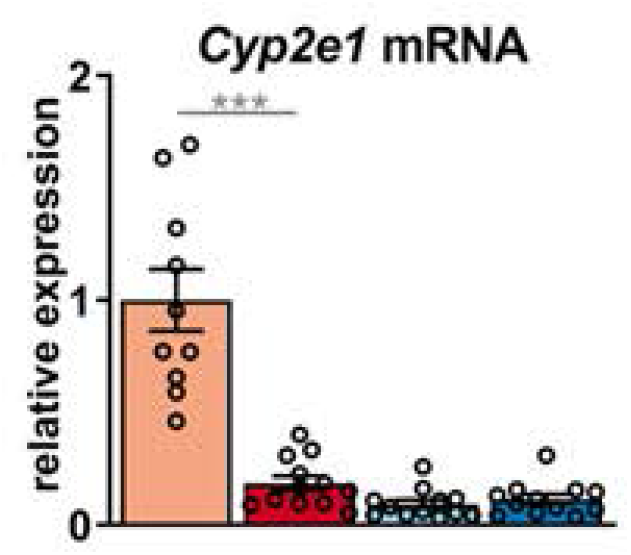
17α-E2 treatment does not modulate Cyp2e1 expression. Liver *Cyp2e1* mRNA at the conclusion of the 8-week intervention [n=10-12/group]. All data are shown as mean ± SEM and were analyzed by one-way ANOVA with Tukey post-hoc testing. We did not indicate statistical differences between vehicle and 17α-E2 treatment groups (preventive and therapeutic), or between 17α-E2 treatment groups (preventive and therapeutic), for purposes of visual clarity. ***p<0.005.

**Supplemental Figure 2:**
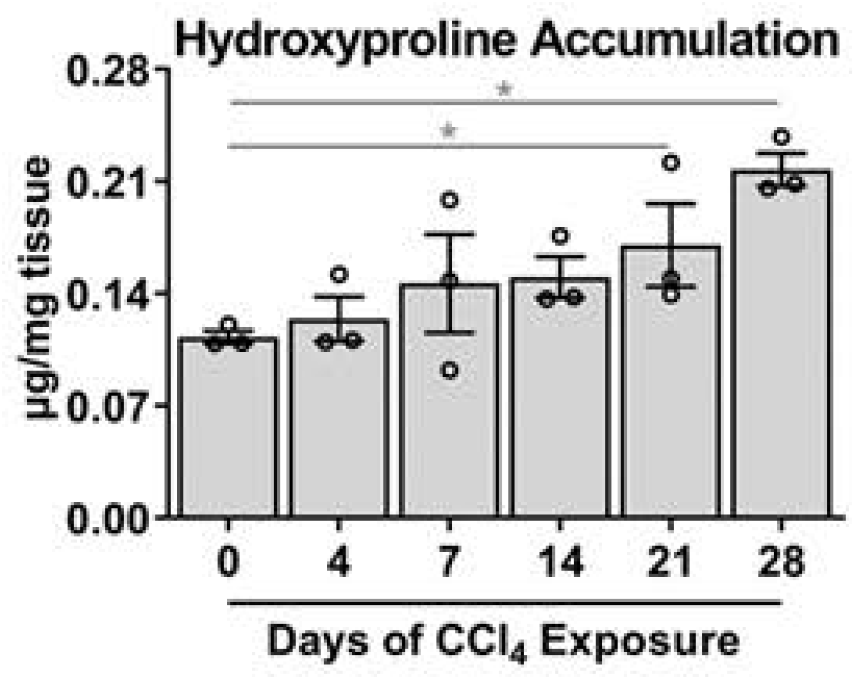
CCl_4_ administration steadily increases liver collagen accumulation over a 4-week period. Liver hydroxyproline accumulation at baseline (day 0) and after 4, 7, 14, 21, and 28 days of CCl_4_ administration [n=3/timepoint]. All data are shown as mean ± SEM and were analyzed by one-way ANOVA with Tukey post-hoc testing. *p<0.05.

**Supplemental Figure 3:**
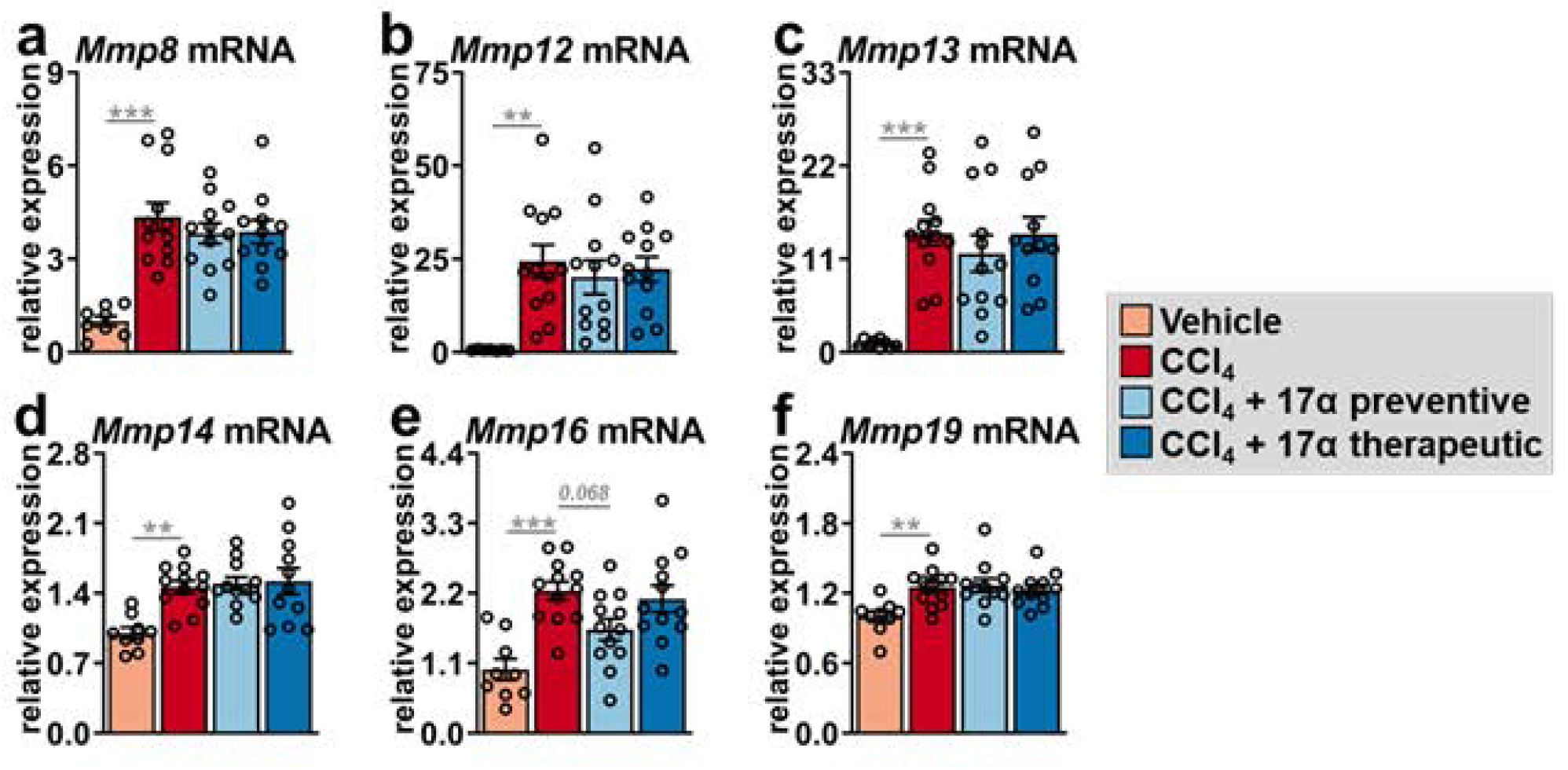
17α-E2 treatment does not alter the expression of MMPs shown to modulate collagen unwinding. Liver (a) *Mmp8* mRNA [n=8-12/group], (b) *Mmp12* mRNA [n=9-12/group], (c) *Mmp13* mRNA [n=9-12/group], (d) *Mmp14* mRNA [n=9-12/group], (e) *Mmp16* mRNA [n=9-12/group], and (f) *Mmp19* mRNA [n=9-12/group] at the conclusion of the 8-week intervention. All data are shown as mean ± SEM and were analyzed by one-way ANOVA with Tukey post-hoc testing. We did not indicate statistical differences between vehicle and 17α-E2 treatment groups (preventive and therapeutic), or between 17α-E2 treatment groups (preventive and therapeutic), for purposes of visual clarity. **p< 0.01, ***p<0.005.

**Supplemental Figure 4:**
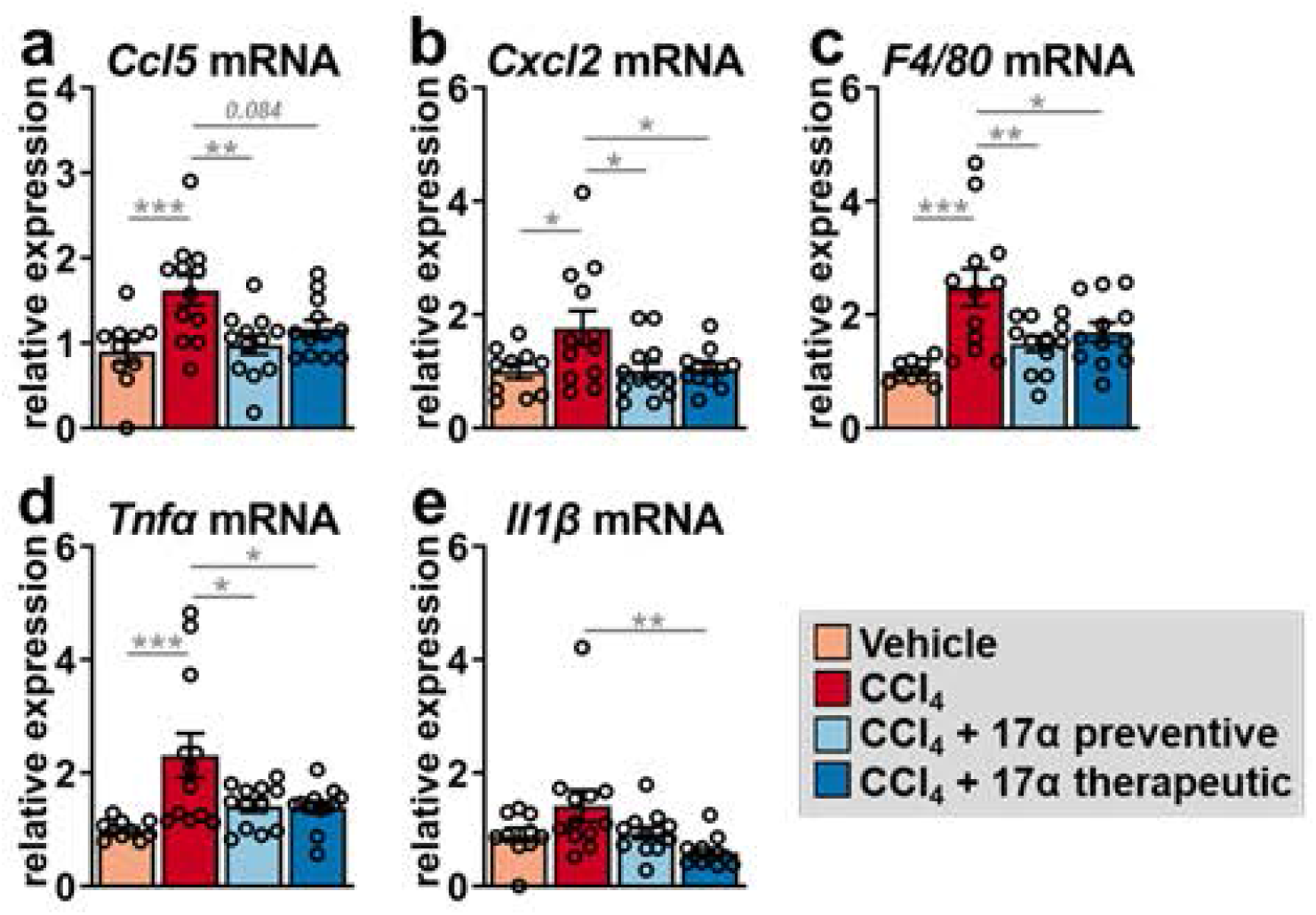
17α-E2 treatment suppresses markers of immune cell recruitment and pro-inflammatory responses in liver. Liver (a) *Ccl5* mRNA [n=9-12/group], (b) *Cxcl2* mRNA [n=10-12/group], (c) *F4/80* mRNA [n=9-12/group], (d) *Tnfα* mRNA [n=10-12/group], and (e) *Il1β* mRNA [n=9-12/group] at the conclusion of the 8-week intervention. All data are shown as mean ± SEM and were analyzed by one-way ANOVA with Tukey post-hoc testing. We did not indicate statistical differences between vehicle and 17α-E2 treatment groups (preventive and therapeutic), or between 17α-E2 treatment groups (preventive and therapeutic), for purposes of visual clarity. *p<0.05, **p< 0.01, ***p<0.005.

